# Development of oocytes in a post-spawning Japanese eel *Anguilla japonica* Temminck & Schlegel in captivity

**DOI:** 10.1101/494435

**Authors:** Satoru Tanaka, Akihiro Okamura, Naomi Mikawa, Yoshiaki Yamada, Noriyuki Horie, Tomoko Utoh, Huan Zhang, Hideo P. Oka, Katsumi Tsukamoto

**Affiliations:** IRAGO Institute Co., Ltd., Tahara, Aichi 441-3605, Japan; Department of Marine Sciences, University of Connecticut, CT 06340, USA; Hamamatsu, Shizuoka 431-0102, Japan; Department of Aquatic Bioscience, Graduate School of Agricultural and Life Sciences, The University of Tokyo, 1-1-1 Yayoi, Bunkyo-ku, Tokyo, 113-8657 Japan

**Keywords:** semelparity, anguillid eel, artificial maturation, spontaneous maturation, water temperature

## Abstract

It is generally believed that parent freshwater eels (*Anguilla* sp.) die soon after spawning on the assumption that eels are a semelparous (or monocyclic) fish (spawn once at the last stage of life) like Pacific salmonids. However, we observed for the first time a post-spawning female *Anguilla japonica* again possessed developing oocytes reaching the final maturation stage in captivity five months after the last spawning even without hormonal treatment. Here we describe information on this female about its biological characteristics including gonadal histology and endocrine profiles. The data suggest that lowering water temperature for a period of time is one of the important factors influencing spontaneous gonadal development in this specimen. We also discuss the possibility of induced multiple spawning of this species in captivity.

---

Freshwater eels (*Anguilla* sp.) are a food resource of worldwide importance, but they are becoming globally endangered, especially in three temperate species European eel *Anguilla anguilla* L., American eel *A. rostrata* (Lesueur, 1817) and *A. japonica* Temminck & Schlegel (Casselman 2003; Dekker 2009; Tsukamoto, Aoyama & Miller 2009). To conserve these eels, techniques for artificial maturation of parent eels by hormonal administration and production of glass eels as seedlings for aquaculture industries have been intensively studied over the past four decades (Yamamoto & Yamauchi 1974; Tanaka 2003; Pedersen 2003; Oliveira & Hable 2010).

Usually, artificially matured parent eels by hormonal treatments are ordinarily discarded after spawning on the assumption that eels are a semelparous (or monocyclic) fish (spawn once at the last stage of life) like Pacific salmonids (*Oncorhynchus* sp.) (Wallace & Selman 1981; Murua & Saborido-Rey 2003). However, we observed for the first time a post-spawning female *A. japonica* again possessed developing oocytes reaching the final maturation stage in captivity five months after the last spawning even without hormonal treatment. Here we describe information on this female about its biological characteristics including gonadal histology and endocrine profiles, and we discuss the possibility of induced multiple spawning of this species in captivity.

A female silver eel of 67 cm in total length (TL) and 445 g in body weight (BW) was captured by a set net in Mikawa Bay (34°N, 137°E) on 19 November 1996 (day 0). Estimated age by the number of the otolith annuli was eight. After acclimatization to an indoor tank (10,000 l) of sea water (34.5 practical salinity unit (PSU)) maintained at 18-20 °C for 2 weeks (day 13), the female was artificially induced to maturation by repeated injection of salmon pituitary extract (SPE, 20 mg kg BW^−1^, once a week) for 6 weeks (Ohta, Kagawa, Tanaka, Okuzawa, Iinuma & Hirose 1997; Kagawa 2003) (Fig. 1). Some males (200–300 g BW), which were purchased from a commercial dealer, were also injected repeatedly with human chorionic gonadotropin (Sankyo, Tokyo, Japan) at a dose of 200 IU per fish once a week at 20 °C until full maturation (Ohta et al. 1997). Then the female was finally injected with luteinizing hormone-releasing hormone (LH-RH, 1 mg kg BW^−1^) (Satoh, Yamamori & Hibiya 1992), and set in the 10,000-L tank together with some matured males (day 58). Two days later (day 60), the female spontaneously released approximately 140,000 eggs based on calculation by the subtraction of BW after spawning (406 g) from that before spawning (493 g). Fertilization rate were about 38.9% and the diameters of fertilized eggs was 1.55-1.65 mm, similar to *A. japonica* fertilized eggs collected at the southern part of the Mariana Ridge (1.49–1.71 mm; n = 12) (Yoshinaga, Miller, Yokouchi, Otake, Kimura, Aoyama, Watanabe, Shinoda, Oya, Miyazaki, Zenimoto, Sudo, Takahashi, Ahn, Manabe, Hagihara, Morioka, Itakura, Machida, Ban, Shiozaki, Ai & Tsukamoto 2011). About 40,000 larvae hatched and survived until 10 days after hatching without feeding. The number of eggs released by the female was considerably smaller than the estimated fecundity of this species ranged from 5 to 10 million eggs for female silver eels (35.7 to 92.4 cm TL) captured at Mikawa Bay (Matsui 1972), suggesting that almost all oocytes possibly remained to be spawned in the abdomen. However, the abdomen gradually shrunk, and the genital pore was completely closed soon after spawning.

**Figure 1.**
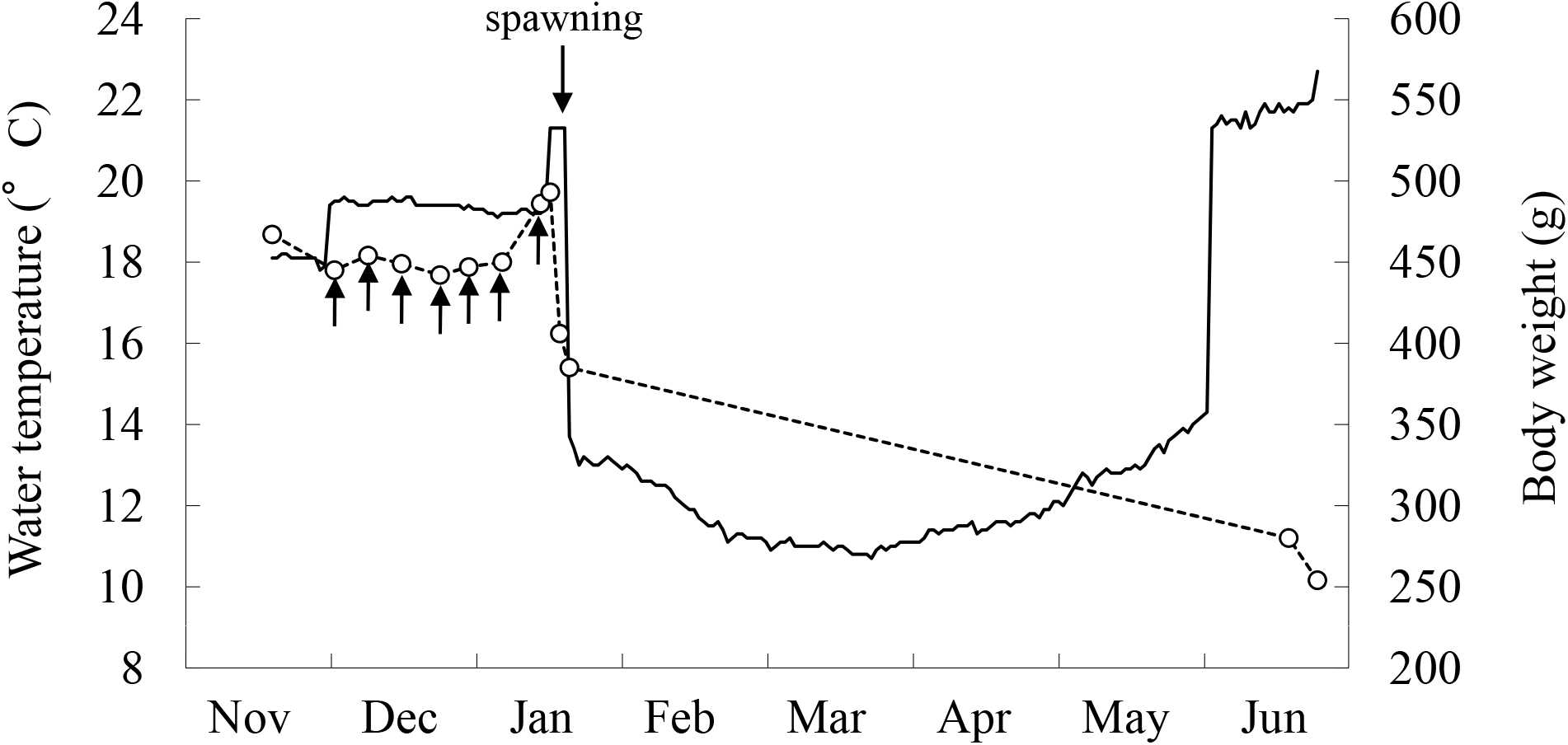
Changes in body weight (dotted line) of the *Anguilla japonica* female, which was once induced spawning and then reared for about 5 months, and water temperature experienced by the female (solid line). Arrows indicate the timings of hormonal injection.

Two days after spawning (day 62), the female was moved to an outdoor tank (2,000 l) of seawater (31.5 PSU) and kept for 134 days without diets (Fig. 1). The water temperature of the tank was once decreased to 10.7°C on day 122 and increased gradually to 14.3 °C until day 196. On day 196, we found that the body of the specimen was markedly thinned compared to the initial body, but the abdomen was somewhat swelling and pliable despite the long-term starvation. Therefore, in anticipation of second spawning, the female was again moved to the indoor tank of water temperature at 22°C and kept for 22 days (Fig. 1). However, spawning did not occur, so we dissected the female after heavy anesthetization on day 218 to observe the gonads macroscopically and microscopically according to the procedure described in Yamamoto, Omori & Yamauchi (1974). The BW of the specimen was decreased to 245 g, about 60% of the BW just after spawning, on day 218 (Fig. 1). Although we have no detailed data on during the starvation, there is the possibility that the female once attained a weight of less than 245 g and the weight increased again as the result of oocyte hydration.

The female had apparently advanced ovaries with gonad-somatic index (GSI = 100 GW BW^−1^; GW: gonad weight) value of 15.4 compared with those of migrating female eels caught at the coastal waters of Mikawa Bay (range: 0.2-4.3, n=286) (Utoh, Mikawa, Okamura, Yamada, Akazawa, Tanaka, Horie & Oka 2004), whereas their GSI values can reach 17.8-51.4 (n=20) at their spawning by artificial maturation (Okamura, Yamada, Horie, Utoh, Mikawa, Tanaka & Tsukamoto 2008). Part of this high GSI value is probably due to a loss of body weight, however, the thickness of ovaries was enlarged compared with those found in migrating females captured at coastal waters (Utoh *et al*. 2004) and its colour was yellow/white (Fig. 2a) and contained healthy oocytes (*ca*. 700 μm in mean diameter). Histology of the ovaries showed that each oocyte included a migrating germinal vesicle but there were no large lipid droplets visible (Fig. 2b), corresponding to those of the early migratory nucleus stage with oocyte diameters of 600-700 μm found in artificially matured Japanese eels (Yamamoto *et al*. 1974). However, the number and size of yolk globule in these oocytes seemed to be somewhat smaller (Fig. 2b), compared with those often found in artificially matured eels at the same stage (Adachi, Ijiri, Kazeto & Yamauchi 2003). Some of degenerating oocytes were also observed (Fig. 2b). Furthermore, the GSI value of this female is similar to those of post-spawning female Japanese eels caught at the spawning area in the western North Pacific (9.0-13.4, n=3)(Tsukamoto, Chow, Otake, Kurogi, Mochioka, Miller, Aoyama, Kimura, Watanabe, Yoshinaga, Shinoda, Kuroki, Oya, Watanabe, Hata, Ijiri. Kazeto. Nomura, & Tanaka 2011). These post-spawning eels had also relatively advanced ovaries including oocytes at the mid-vitellogenic stage with oocyte diameters of 350-600 μm.

**Figure 2.**
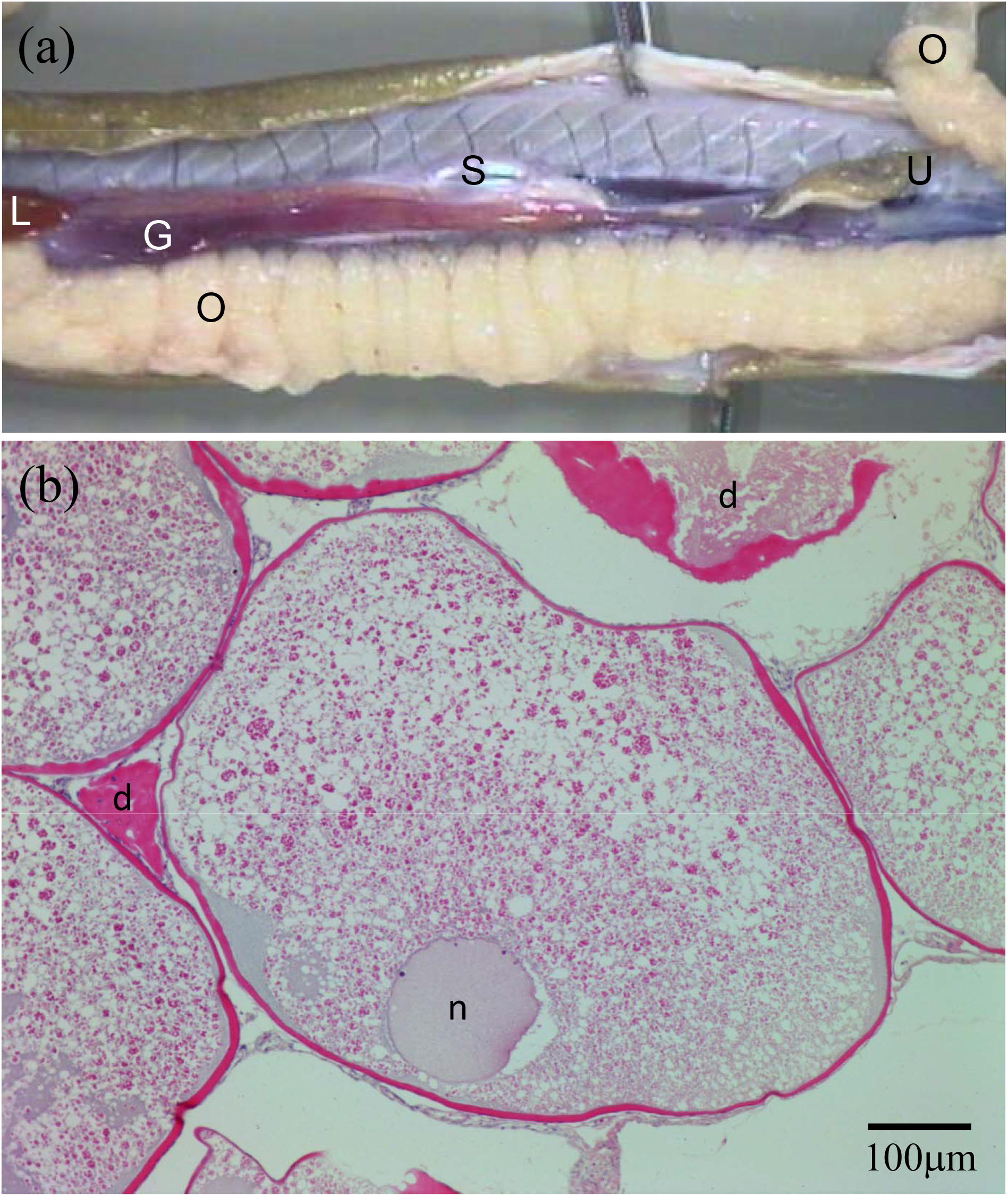
Macroscopic image of ovaries (a) and histological image of oocytes (b) in the *Anguilla japonica* female reared for about 5 months after the last spawning. G, gut; L, liver; O, ovary; S, swim bladder; U, urogenital pore; d, oocyte atresia; n nucleus.

Estradiol-17β (E2) and vitellogenin levels in the serum were also measured by the methods described elsewhere (Asahina, Kambegawa & Higashi 1995; Mikawa, Utoh & Oka, 2001; Utoh, Horie, Mikawa, Okamura, Yamada, Akazawa, Tanaka & Oka 2005). Serum E_2_ levels detected in this female (0.069 ng ml^−1^) was apparently lower than those usually found in artificially matured eels at the migratory nucleus stages (range: 9.4-12.3 ng ml^−1^, n=75) (Matsubara, Lokman, Kazeto, Adachi & Yamauchi 2005). This low E2 levels, however, may be not strange at the post vitellogenic stage, because higher levels of E2 often detected in artificially matured eels at the final maturation stage are considered to be a peculiar feature induced by SPE injections (Chiba, Iwatsuki, Hayami, Hara & Yamauchi 1994; Ijiri, Kazeto, Takeda, Chiba, Adachi & Yamauchi 1995). Generally, serum E2 levels in other teleosts increase in parallel with the progress of vitellogenesis and the decrease at the final maturation stage (Fostier; Jalabert; Billard; Breton & Zohar 1983), suggesting that low E2 levels found in the present female may reflect a normal endocrine profile which appears in wild eels under natural conditions.

Meanwhile, vitellogenin, which is the precursor of yolk protein synthesized in liver in response to E_2_, was detected at 2.52 mg ml^−1^ in the serum, which was a level included in the range found in wild silver eels at the early-to mid-vitellogenic stages caught in Mikawa Bay (range: 0.1-6.5 mg ml^−1^, n=144) (Mikawa *et al*. 2001), but was apparently lower than those found in artificially matured female silver eels (2.5-10 mg ml^−1^ n=20) (Sato, Kawazoe, Suzuki & Aida 2000). This levels of vitellogenin found in the present female are possibly the reminder of circulating vitellogenin, because it should be no longer necessary at this stage.

These reproductive characteristics suggest that this female had been ready to undergo the final maturation phase. Then, why did the female possess developing oocytes five months after the previous spawning? One possibility is that developed oocytes were retained in the female’s abdomen just as it was for five months. This possibility is suggested by the calculation that the number of eggs spawned by the female (140, 000 eggs) was considerably smaller than the usual fecundity of this species (5 to 10 million eggs). However, Changchun, Yujun, Zhengfeng, Yiqiang & Yejin (1980) observed that remnant oocytes in post-spawning Japanese eels were usually absorbed by at least two months after spawning in captivity. Therefore, we think that this possibility is low. Another possibility is that these oocytes newly developed after the previous spawning. The external color of ovaries in the female on day 218 (Fig. 1a) appeared to be fresh and healthy, which were considerably differed from those with a lot of ecchymoses usually found in females just after spawning in captivity (Mikawa, unpublished data), suggesting the de novo development of oocytes in this female. Because the serum levels of the salmon gonadotropin (GTH) injected into eels rapidly decrease to considerable low levels by four days and undetectable levels by 24 days after the last injection (Sato, Kawazoe, Suzuki & Aida 1996; 2000), the influence of residual activity of SPE might be no longer available soon after spawning. If so, we suppose that the development of these oocytes had been controlled by endogenous hormones.

Generally, it has been known that eels do not sexually mature in captivity without exogenous hormonal assistance, because their endogenous hormonal systems do not work under captive conditions, probably due to a lack of appropriate environmental triggers for maturation (Adachi *et al*. 2003; Palstra, Ginneken & van den Thillart 2009). Perhaps, relatively lower water temperature for five months experienced by the present female is likely one of important factors influencing its gonadal development. Tracking experiment of silver *A. anguilla* revealed that they undergo diel vertical migration during the spawning migration between 300 and 500 m in depth where temperatures were lower than 13°C (Tesch 1989). Another tracking study showed that silver *A. dieffenbachii* Gray occasionally dove to around 6°C (Jellyman & Tsukamoto 2010). A laboratory experiment also indicated that a variable thermal regime that increased from lower (10 °C) to higher temperatures (14 and 17 °C) induced a faster gonadal development, higher gonadotropin β-subunits (fshβ and lhβ) expression and higher serum E2 levels during vitellogenesis in European female eels injected with carp pituitary (Pérez, Peñaranda, Dufour, Baloche, Palstra, van den Thillart & Asturiano 2011). In a marine eel, *Conger myriaster* (Brevoort), gonadal development completes without hormonal treatment in captivity at 6°C and they spawn several days after an abrupt temperature elevation to 10 °C (Utoh & Horie 2011). Therefore, it is possible that relatively lower water temperatures (11-14°C) for five months stimulated pituitary hormonal axis and induced spontaneous oocyte development. The abrupt elevation of water temperature from 14 to 22°C (Fig. 1), however, might be unfavorable for the female reaching final maturation phase. Satoh *et al*. (1992) and Dou, Yamada, Okamura, Shinoda, Tanaka & Tsukamoto (2007) showed that spontaneous spawning of artificial matured *A. japonica* is efficiently induced by an elevation of water temperature from 18 or 20 to 22°C. However, the present female did not spawn after the temperature elevation from 14 to 22°C. It is possible that the timing or width of temperature elevation was not adequate for the female in the present study. These inadequate temperature conditions during the final maturation phase were likely reflected in a relatively small number of yolk globules in oocytes and some of degenerating oocytes found in this female (Fig. 2b).

Previously, Changchun *et al*. (1980) also observed histologically a number of newly developed oocytes at the pre-vitellogenic stage in post-spawning Japanese eels (n=23) two years after spawning and they did not die but ingested diets and regained their BW after spawning. Horie *et al*. (unpublished data) observed that a number of post-spawning *A. japonica* females repeatedly spawned (at least 3 times) in captivity two or three weeks following previous spawning events when they were continually received hormonal administration. These evidences suggest that eel ovary is possibly polycyclic and the type of ovarian development of organization in this species might be ‘group-synchronous’ rather than ‘synchronous’ (Wallace & Selman 1981; Murua & Saborido-Rey 2003).

In conclusion, the post-spawning female Japanese eel possessed almost matured oocytes five months after spawning, suggesting that captive eels have the potential of multiple spawning. In addition, lower water temperatures for a period of time may be one of the important factors influencing spontaneous gonadal development in this species, which may lead to establishing a technique to obtain efficiently eggs or to induce naturally eel maturation without exogenous hormones.

